# “Organ-in-a-column” coupled on-line with liquid chromatography-mass spectrometry

**DOI:** 10.1101/2020.09.08.282756

**Authors:** Stian Kogler, Aleksandra Aizenshtadt, Sean Harrison, Frøydis Sved Skottvoll, Henriette Engen Berg, Shadab Abadpour, Hanne Scholz, Gareth Sullivan, Bernd Thiede, Elsa Lundanes, Inger Lise Bogen, Stefan Krauss, Hanne Røberg-Larsen, Steven Ray Wilson

**Author notes:** Corresponding author: Steven Ray Wilson. Full address: Department of Chemistry, University of Oslo, Post Box 1033, Blindern, NO-0315 Oslo, Norway.

## Abstract

Organoids, i.e. laboratory-grown organ models developed from stem cells, are emerging tools for studying organ physiology, disease modeling and drug development. On-line analysis of organoids with mass spectrometry would provide analytical versatility and automation. To achieve these features with robust hardware, we have loaded liquid chromatography column housings with induced pluripotent stem cell (iPSC) derived liver organoids and coupled the “organ-in-a-column” units on-line with liquid chromatography-mass spectrometry (LC-MS). Liver organoids were co-loaded with glass beads to achieve an even distribution of organoids throughout the column while preventing clogging. The liver organoids were interrogated “on column” with heroin, followed by on-line monitoring of the drug’s phase 1 metabolism. Enzymatic metabolism of heroin produced in the “organ-in-a-column” units was detected and monitored using a triple quadrupole MS instrument, serving as a proof-of-concept for on-line coupling of liver organoids and mass spectrometry. Taken together, the technology allows direct integration of liver organoids with LC-MS, allowing selective and automated tracking of drug metabolism over time.

---

Drug discovery and development is an extremely costly process, and the number of new drugs reaching the market per billion dollars spent on research and development is consistently low. Moreover, the efficacy/toxicity of drugs can vary significantly in target patients, calling for personalized drug testing [1]. Key bottlenecks of efficient drug development include the limited predictive value of traditional cell cultures and animal models for human drug metabolism, and personalization of the model systems [2]. Hence, novel technologies for predicting personalized drug metabolism are being explored.

Organoids, here broadly defined as *in vitro* 3D models that exhibit features of the mature organ in question, are rapidly emerging as powerful tools for drug discovery and personalized testing. Organoids can be readily derived from stem cells and carry the potential for serving as relevant and personalized testing materials [3]. Typically, organoids are 200-500 μm in size, consisting of organ-specific cell types [4]. For example, liver organoids can consist of hepatocytes, hepatic stellate cells, endothelial cells, and cholangiocytes, and can be used as a tool for assessing aspects of drug metabolism and toxicity [5, 6].

For measuring small molecules such as drugs and their metabolites, liquid chromatography-mass spectrometry (LC-MS) is a method of choice in analytical chemistry. Mass spectrometric analysis of organoids has been performed indirectly (“off-line”), i.e. samples are collected and handled semi-manually prior to MS [7–10]. Off-line handling can be time consuming and prone to variations, depending on the method, sample size, and analyte stability. Direct on-line MS analysis of organoids would potentially offer the advantage of increased automation and possibly improved throughput. Verpoorte and co-workers have previously combined organs/organ models, chip microfluidics and separation science, for studying metabolism of liver slices [11] and pharmacology in a gut-on-chip [12]. We have recently coupled liver organoids with sample preparation techniques such as electromembrane extraction (EME) [13], which we also found to be compatible with on-line coupling of organoid-containing chips to LC-MS for studying organoid drug metabolism [14]. However, EME requires an electrical current driven transfer of metabolites through an oil membrane into an MS-compatible solution which can potentially limit the spectrum of analytes that can be analyzed [14]. Additionally, a key challenge for coupling chips with MS is ensuring practical and robust connections and standardization. Therefore, we have explored placing liver organoids directly into standardized/commercial tubing and connectors of liquid chromatography (LC), perhaps the most applied fluidics platform in analytical chemistry. Specifically, LC column housings were loaded with organoids generated from iPSC-derived hepatocyte-like cells (iHLC organoids) and sandwiched between an upstream drug delivery system and a downstream connector to a traditional LC-MS setup. The system, termed “organ-in-a-column” allowed in-column cultivation of liver organoids for an extended period, “on-line” exposure to drugs and monitoring of drug metabolism using mass spectrometry.

## Experimental

### Consumables and basic hardware

Stainless steel (SS) unions, reducing unions (1/16” to 1/32”), SS ferrules and nuts (all for 1/16” tubing and for 1/32” tubing), SS tubing (1/32” outer diameter (OD), 0.020” inner diameter (ID) and 0.005” ID), perfluoroalkoxy alkane (PFA) tubing (1/16” OD, 0.75 mm ID) and 1/16” SS screens (2 μm pores) were purchased from VICI Valco (Schenkon, Switzerland). SST Vipers (130 μm x 650 mm) were purchased from Thermo Fisher Scientific (Waltham, MA, USA). A chromatographic column (1 mm x 5 cm) packed with Kromasil C4 (3.5 μm particles, 100 Å pore size) was purchased from Teknolab (Ski, Norway). Luer lock syringes (10-20 mL) were purchased from B. Braun Melsungen AG (Hessen, Germany). Acid-washed glass beads (150 – 212 μm) were purchased from Sigma Aldrich (St. Louis, MO, USA).

### Reagents and solutions

Formic acid (FA, ≥ 98%) was purchased from Merck (Darmstadt, Germany). Water (LC-MS grade) and acetonitrile (ACN, LC-MS grade) were purchased from VWR International (Oslo, Norway). Tough 1500 3D-printer resin was purchased from Formlabs Inc. (Somerville, MA, USA). For liquid chromatography, mobile phase (MP) reservoir A contained 0.1% FA in HPLC water (v/v). MP reservoir B contained ACN/HPLC water/FA (90/10/0.1%, v/v/v).

Heroin HCl, 6-acetyl morphine (6-AM) HCl, and morphine were obtained from Lipomed AG (Arlesheim, Switzerland). Heroin-d9, 6-AM-d6, and morphine-d3 (used for heroin stability experiments only) were purchased from Cerilliant (Austin, TX, USA). Fetal bovine serum-free medium (William’s E medium, supplemented with 0.1 μM dexamethasone and 1% insulin-transferrin-selenium mix) and L15 base medium (prepared according to [15]) is hereafter referred to as organoid medium.

### Organoids

iHLC organoids originating from 3 cell lines (iHLC-1 = WTC-11, iHLC-2 = WTSIi013-A, and iHLC-3 = WTSIi028-A, Wellcome Trust Sanger Institute) were differentiated toward liver organoids using a modification of the protocol by Lee et al. [16]. iPSC line AG27 was differentiated to form liver organoids containing induced hepatocyte like cells (iHLCs) as described by Harrison et al. [10] and was used in initial experiments (and figures 5 and SI1). Cryopreserved primary human hepatocytes (PHH, Gibco, catalogue no. HMCPMS, lot HU8287) were thawed in the Hepatocytes thaw media (Gibco, catalogue no. CM7500) according to manufacturer’s protocol. Uniform PHH spheroids were created by aggregation in house-made agarose microwells as described before [17] and cultured in Williams E medium (Thermo Fisher Scientific, catalogue no. A1217601) supplemented with 0,5 % FBS (Thermo Fisher Scientific, catalogue no. 41400045), 2 mM L-glutamine (Thermo Fisher Scientific, catalogue no. 35050038), 10 μg/ml insulin, 5.5 μ g/ml transferrin, 6.7 ng/ml sodium selenite (Thermo Fisher Scientific, catalogue no. 41400045) and 0.1 μM dexamethasone (Sigma Aldrich, catalogue no. D4902).

### Instruments and advanced hardware

The Dionex UltiMate 3000 UHPLC system and the TSQ Vantage MS with the HESI-II ion source were purchased from Thermo Fisher Scientific. The syringe pump (AL-1000) was bought from World Precision Instruments (Sarasota, FL, USA). A 2-position 10-port valve (for 1/32”, C82X-6670ED) was purchased from VICI Valco. A SUB Aqua 5 Plus water bath was purchased from Grant Instruments (Cambridge, UK). The Form 3B 3D-printer and wash and cure station were purchased from Formlabs Inc (Somerville, MA, USA). A PST-BPH-15 column heater was purchased from MS Wil (Aarle-Rixtel, the Netherlands). The refrigerated circulating water bath was purchased from Haake (Berlin, Germany).

### Heroin stability testing

Stability testing of heroin was performed by incubating solutions of L15 base medium, serum-free organoid medium, and type 1 water. The incubation was performed at 4 °C and 37 °C. For each solution, 100 μL freshly made 1 mM heroin (in 0.9 % NaCl) or 0.9 % NaCl (control) and 900 μL organoid medium were mixed. At 0, 24, 48, and 120 hours, 100 μL samples were collected from all solutions. To precipitate proteins, 10 μL of 1.1 M FA was added to each sample, followed by vortexing and 2 min centrifugation at 14500 g. 50 μL of the resulting supernatant was transferred to a new vial and diluted to 1 mL with type 1 water. Samples were stored at −80°C prior to analysis. For these experiments, the determination of heroin, 6-AM, and morphine were performed using UHPLC-MS as previously described by Skottvoll *et al*. [13].

### 3D printed syringe cooler

A 3D-printed double-wall syringe cooler was printed and fitted directly to the syringe cylinder **(Figure 1A).** 3D-printed syringe coolers were designed using SOLIDWORKS CAD-software (3DS, Paris, France). Syringe coolers were printed in Tough 1500 resin with the Form 3B 3D-printer. A cross-section of the double-walled design is shown in **Figure 1B.** Wall thickness in the main body was 1 mm. The flow through part of the body was 5 mm thick, and the inner diameter was adapted to fit the individual syringe. Note that dimensions of e. g. a 3 mL syringe varies greatly between different manufacturers. Cold water (4 °C) was pumped through the cooler with a refrigerated circulating water bath **(Figure 1C).** To ensure a cold stable temperature from the start, the water bath, organoid medium, and heroin solutions were cooled prior to the start of an experiment. Due to the low flow rate (15 μL/h), and a small ID of the tubing (0.75 mm), the medium is quickly heated to physiological temperature upon introduction to the “organ-in-a-column” which is kept at 37 °C in a column heater.

**Fig 1.**
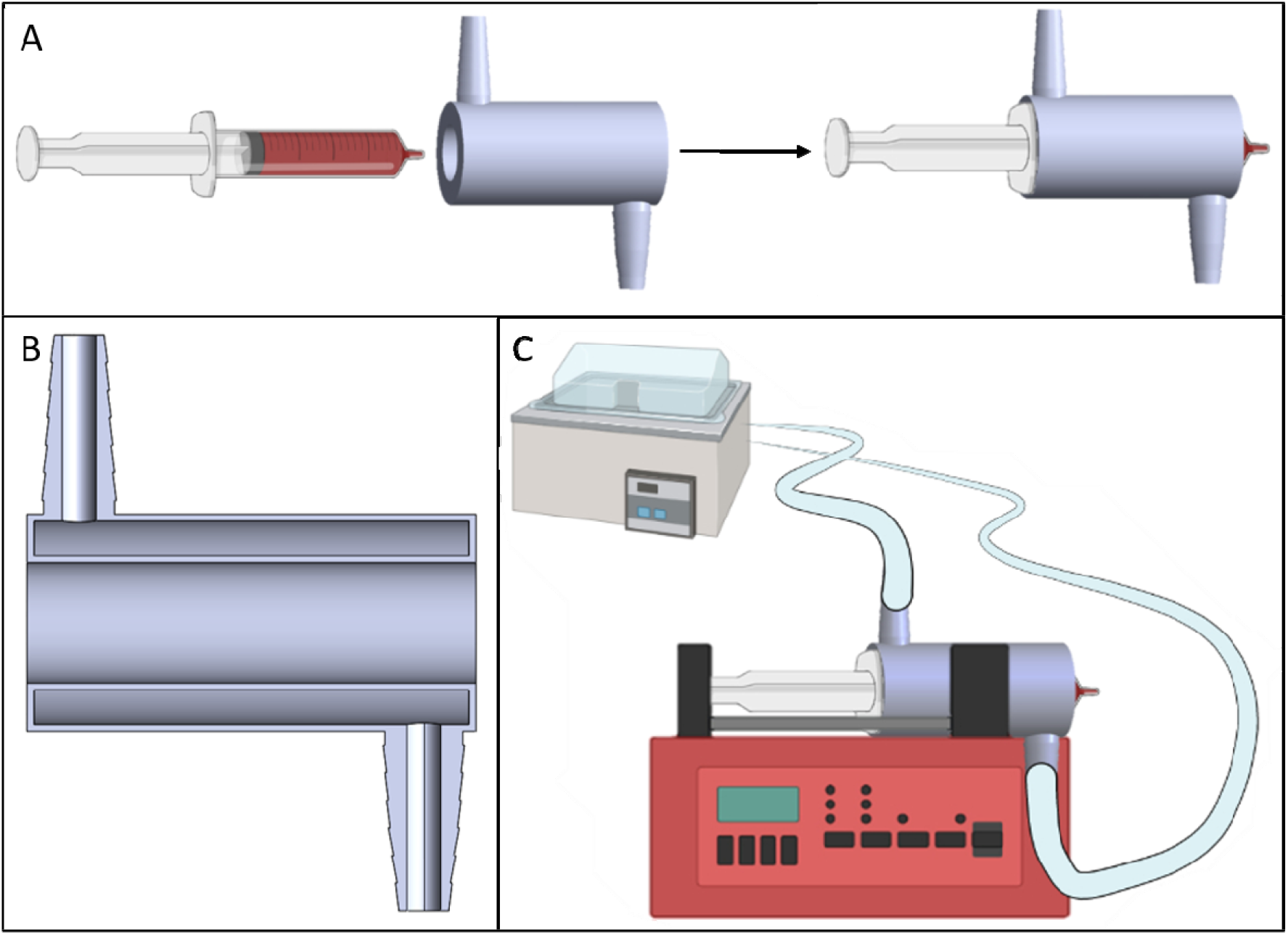
A: The 3D-printed syringe cooler was tailored to fit the syringe cylinder, ensuring a snug fit. 10 mL Luer lock syringes from B. Braun Omnifix were used. B: Cross section of the syringe cooler’s double-walled design. Wall thickness was 1 mm. The water chamber was 5 mm thick. C: The syringe cooler was connected to a refrigerated circulation bath from Haake.

### “Organ-in-a-column”

For fabrication of the column housing, a 10 cm long piece of PFTE/PFA tubing (1/16” OD, 0.75 mm ID) was cut and assembled with nuts and SS ferrules. To one end of the tube, a union with a 1 μm SS screen (VICI Valco) was connected. Organoid medium containing approximately 50 iHLC organoids (size range 100 μm - 200 μm) was then transferred to a 3 mL Luer lock syringe. Two spatulas of acid washed glass beads, containing approximately 45 mg of beads, were subsequently added. Through gentle shaking, beads and organoids were mixed in the syringe. The syringe was then connected to the open end of the column. By pressing the contents of the syringe through the empty column housing, the organoid column was finalized. Once the entire contents of the syringe were passed through the column, the inlet was fitted with a SS screen and a union. A schematic of an “organ-in-a-column” is shown in **Figure 2.** For idle conditions, a new syringe was filled with fresh organoid medium, connected to the “organ-in-a-column”, and placed in a syringe pump. The pump was set to 15.0 μL/h. After filling, the “organ-in-a-column” was kept in the column heater at 37 °C.

**Figure 2.**
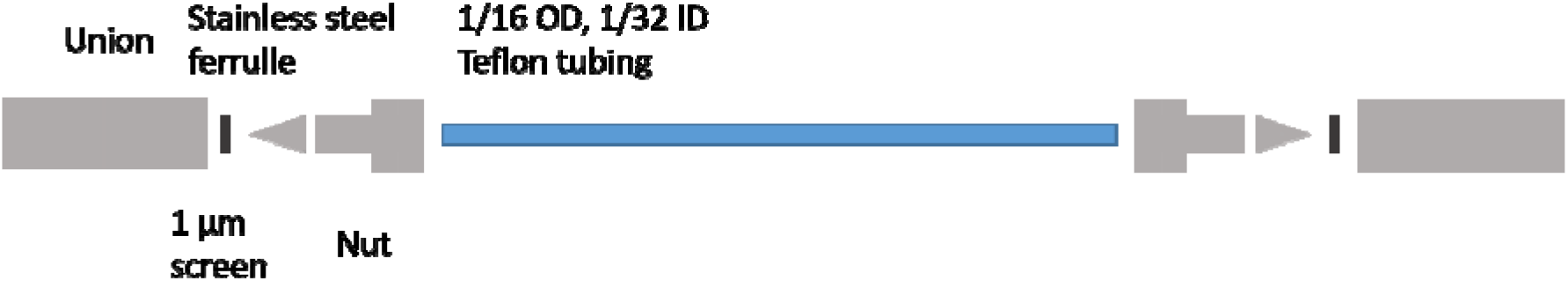
Column housing for “organ-in-a-column”.

### “Organ-in-a-column” coupled with liquid chromatography-mass spectrometry

The pump/syringe system described above was connected to LC-MS instrumentation. Eluate from the “organ-in-a-column” was transported to a valve system for fractionation. The valve system contained two sample loops (5 μL), which were filled sequentially. As one loop was being filled, the content of the other loop was pumped to a 5 cm x 1 mm C4 LC column for separation prior to MS detection. The 5 μL loops were overfilled with an additional 2.5 μL to ensure that any LC solvent left in the loop was flushed off and to accommodate possible small fluctuations of syringe pump flow rate. See **Figure 3** for a schematic of the setup. For studying heroin metabolism, a fresh stock solution of 1 mM heroin HCl in 0.9% NaCl was prepared prior to each experiment and diluted with organoid medium to 10 μM. The organoid medium/heroin-solutions were pumped at 15.0 μL/h, with fractionation every 30 minutes. This allowed for 5 μL injections onto the LC-MS system, with an overfilling of a factor 1.5 (= 7.5 μL, delivered in 30 minutes at the 15 μL/h flow rate) to ensure proper loop filling. See Table 1–3 for LC-MS conditions (C4 1 mm ID separation column, flow rate 50 μL/min).

**Figure 3.**
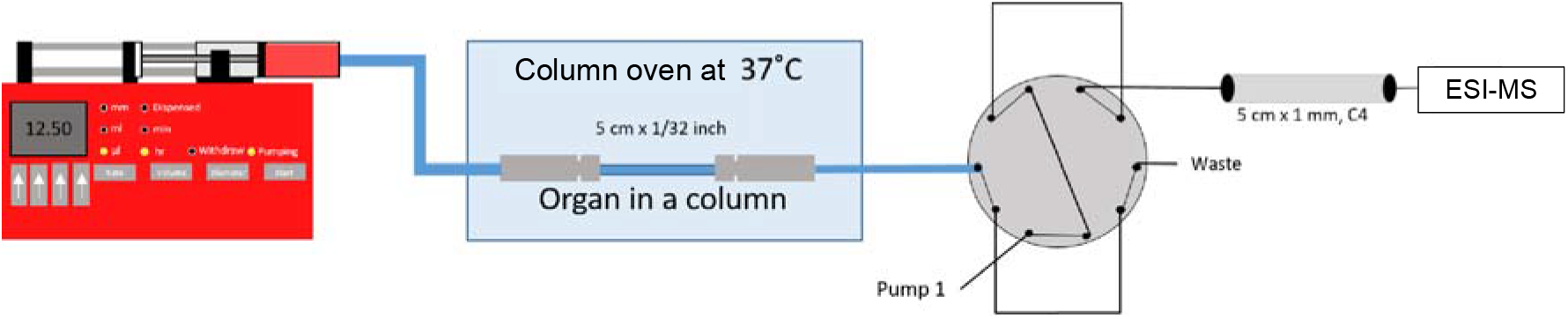
Illustration of “organ-in-a-column” coupled on-line with liquid chromatography-mass spectrometry. See the experimental section for more details.

**Table 1.** LC-gradient for on-line studies of “organ-in-a-column” metabolism of heroin.

| Time, min | Flow, $\mu\text{L}/\text{min}$ | Mobile phase, %B | Purpose |
| --- | --- | --- | --- |
| 0-2 | 50 | 3 | Separation |
| 2-10 | 50 | 3-20 |  |
| 10-20 | 50 | 80 | Wash |
| 20-30 | 50 | 3 | Re-equilibration |

**Table 2.** Multiple reaction monitoring (MRM) parameters used for the detection of heroin and its phase 1 metabolites.

| Analyte | Parent ion, $m/z$ | Product ion, $m/z$ | Collision energy, eV | S-lens value |
| --- | --- | --- | --- | --- |
| Heroin | 370.15 | 286.04 | 42 | 104 |
|  |  | 210.96 | 59 | 104 |
| 6-AM | 328.13 | 210.96 | 6 | 72 |
|  |  | 180.97 | 37 | 72 |
| Morphine | 286.14 | 200.99 | 5 | 88 |
|  |  | 184.91 | 48 | 88 |

**Table 3.** General MS parameters for detection of heroin, 6-AM and morphine.

| Parameter | Value |
| --- | --- |
| Capillary temperature | 300.0 °C |
| Vaporizer temperature | 200.0 °C |
| Sheath gas pressure | 20.0 |
| Ion Sweep gas pressure | 0.0 |
| Auxillary air flow | 5.0 |
| Spray voltage | Pos: 3000.0 V Neg: 0.0 V |
| Collision gas pressure | 1.0 mTorr |

### Proteomic analysis of iHLCs and PHHs using liguid chromatography-tandem mass spectrometry

Heroin treated (harvested after 24 hours) iHLC organoids generated from 3 cell lines (iHLC-1 = WTC-11 (WiCell), iHLC-2 = WTSIi013-A, and iHLC-3 = WTSIi028-A, Wellcome Trust Sanger Institute) and 3D spheroids generated from cryopreserved primary human hepatocytes (PHHs, Gibco, lot HU8287) were prepared in 2 replicates in addition to controls (untreated iHLCs and PHHs, n=1). Pelleted iHLC organoids and PHH spheroids were prepared by Easy Extraction and Digestion (SPEED) [18], using dithiothreitol (DTT) and iodoacetamide (IAM) for reduction and alkylation, respectively, and 6 μg trypsin for digestion (overnight at 37 °C). Sample clean-up (after concentrating and reconstitution of samples in 100 μL LC-MS grade water containing 0.25% *(v/v)* heptafluorobutyric acid) was performed using 100 μL Bond Elut C18 solid-phase extraction pipet tips (Agilent, Santa Clara, US) following the protocol of the manufacturer. The eluate was concentrated to dryness and dissolved in 4 μL LC-MS grade water containing 0.1% (v/v) FA.

LC-MS analysis was performed using a timsTOF Pro (Bruker Daltonics, Bremen, Germany) coupled to a nano Elute nanoflow LC system (Bruker Daltonics). Separation was performed with a 25 cm x 75 μm, 1.6 μm, C18, Ion Optics (Fitzroy, Australia) column operated at 50 °C. Mobile phase A and B reservoirs contained LC-MS grade water and acetonitrile, respectively, both containing 0.1% (v/v) formic acid. A linear gradient from 0 - 35% B (54 min) was employed (300 nL/min flow rate). MS acquisition was performed in data-dependent acquisition parallel accumulation-serial fragmentation mode, and an injection volume of 2 μL was employed. Data were searched against the human UniProt database (20,431 entries), with PEAKS X+ software version 10.5 (Bioinformatics Solutions), allowing 1 missed cleavage and a false discovery rate of 1%.

### Measurement of enzyme expression in iHLC organoids and PHH spheroids

RNA was isolated using TRIzol reagent (Thermo Fisher Scientific) according to the manufacturer’s protocol. RNA concentration and purity were analyzed using NanoDrop ND-1000 spectrophotometer (Thermo Fisher Scientific). cDNA was synthesized using High-Capacity cDNA Reverse Transcription Kit (Thermo Fisher Scientific, catalogue no. 4368814). Gene expression analysis was performed using a TaqMan Universal mix on a TaqMan ViiA7 Real Time PCR System. PPIA was used as endogenous control. Level of expression of genes of interest were quantified by ddCt with normalization to control (vehicle-treated organoids).

## Results and Discussion

Liver organoids were generated from four induced pluripotent stem cell (iPSC) cell lines and benchmarked to primary human hepatocytes, grown as spheroids. In support of expected liver functionality, enzymes related to the metabolism of heroin (CES1 and CES2) were analyzed and identified by rtPCR and proteomic analysis in iHLC1-3 liver organoids (see **Supporting Information, SI4-SI5).**

### Liver organoid charging of an LC-column structure

LC column fittings are specifically designed for leakage-free and simple packing, and come in a variety of diameters and lengths, with readily available fittings. Hence, we aimed to investigate whether LC columns could be directly loaded with liver organoids, and whether the liver organoids can be grown for an extended period in the columns without losing viability. LC columns have previously been demonstrated as useful housings for studying biological interactions, e.g. Wiedmer and co-workers’ studies of drug interactions with in-column liposomes [19]. Liver organoids were used for packing LC column housings: iHLC organoids from four different iPSC cell lines.

For liquid chromatography column housings, a micro LC format was chosen (0.75 mm ID and 10 cm length). Both PFA-tubing and PTFE-tubing have been used instead of regular steel column housings as their optical properties allow for visual inspection of organoids *in situ* (see **Figure 4** of stained organoids in the column). To prevent clogging, organoid medium containing liver organoids was supplemented with glass beads (45 mg beads per 10 cm column/50 organoids) prior to packing. After gently mixing the organoid/glass bead containing organoid medium solution, columns were filled manually with a syringe into an open LC column housing. Post-filling, the column was coupled to the upstream and downstream hardware. The addition of the beads kept the liver organoids well-spaced throughout the column, significantly reducing clogging and increasing the robustness of the system.

**Figure 4.**
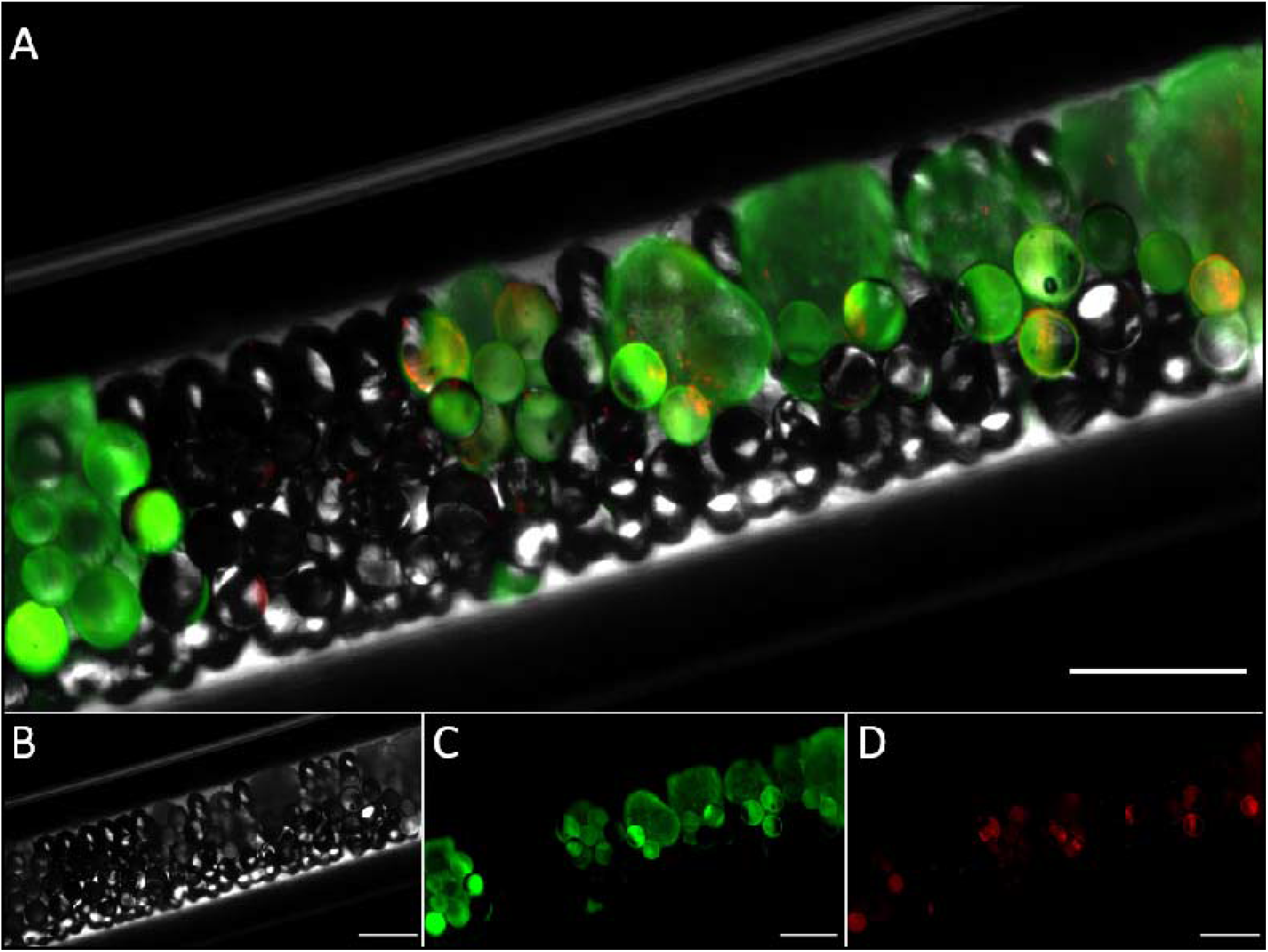
Fluorescence microscopy image of iHLC organoids loaded with glass beads in an “organ-in-a-column”. Viable cells were stained with green fluorescent calcein-AM. Dead cells were stained with propidium iodide (red). Brightfield (B), green (C), and red (D) channels are shown below the merge (A). Scale bar is 600 μm.

After on-line culture and measurements (see below), liver organoids could be readily flushed from the column by removing one of the columns unions and applying mild pressure with a hand-held syringe. Pre-loading and post-flushing inspection of the organoids by microscopy revealed that >80 % of the organoids showed substantial amounts of live staining after 7 days within the perfused column in both the absence and presence of heroin exposure (see **Supporting Information, SI1** for examples).

### Coupling of the liver organoid loaded “organ-in-a-column” to an LC-MS system

The “organ-in-a-column” containing iHLCs was coupled to high-pressure LC through a fractionation valve set-up. LC-MS is generally operated at high pressures e.g. 50-400 bars, which is incompatible with organoid culture. A 2-position 10-port stainless steel valve was used to collect and pump liquid fractions to the LC-MS system, not unlike that used for two-dimensional LC separations [20]. The valve system set-up efficiently isolated the organoids from non-biocompatible solvents and high-pressure of the analysis system (see **Figure 3).**

Organoid medium is complex and can contain considerable amounts of proteins such as albumin. Sample complexity and the presence of proteins can cause unpredictable chromatographic performance. A short-chained butyl (C4) stationary phase (considered relatively compatible with proteins) allowed for repeatable chromatography of organoid medium spiked with model substance heroin/metabolites at this stage of the project (see **Figure 5A).** Mass spectrometric detection was performed in multiple reaction monitoring (MRM) mode, which allowed highly selective and sensitive detection of small molecules such as heroin and its metabolites 6-AM and morphine **(Figure 5B,** from initial experiments with AG27-derived organoids). The mobile phase composition was also a key parameter regarding robustness; methanol as an organic modifier was associated with column clogging and poor performance when chromatographing the organoid medium, while acetonitrile provided significantly improved performance. See **Figure 5C** for illustration of the retention time repeatability for the system.

**Figure 5.**
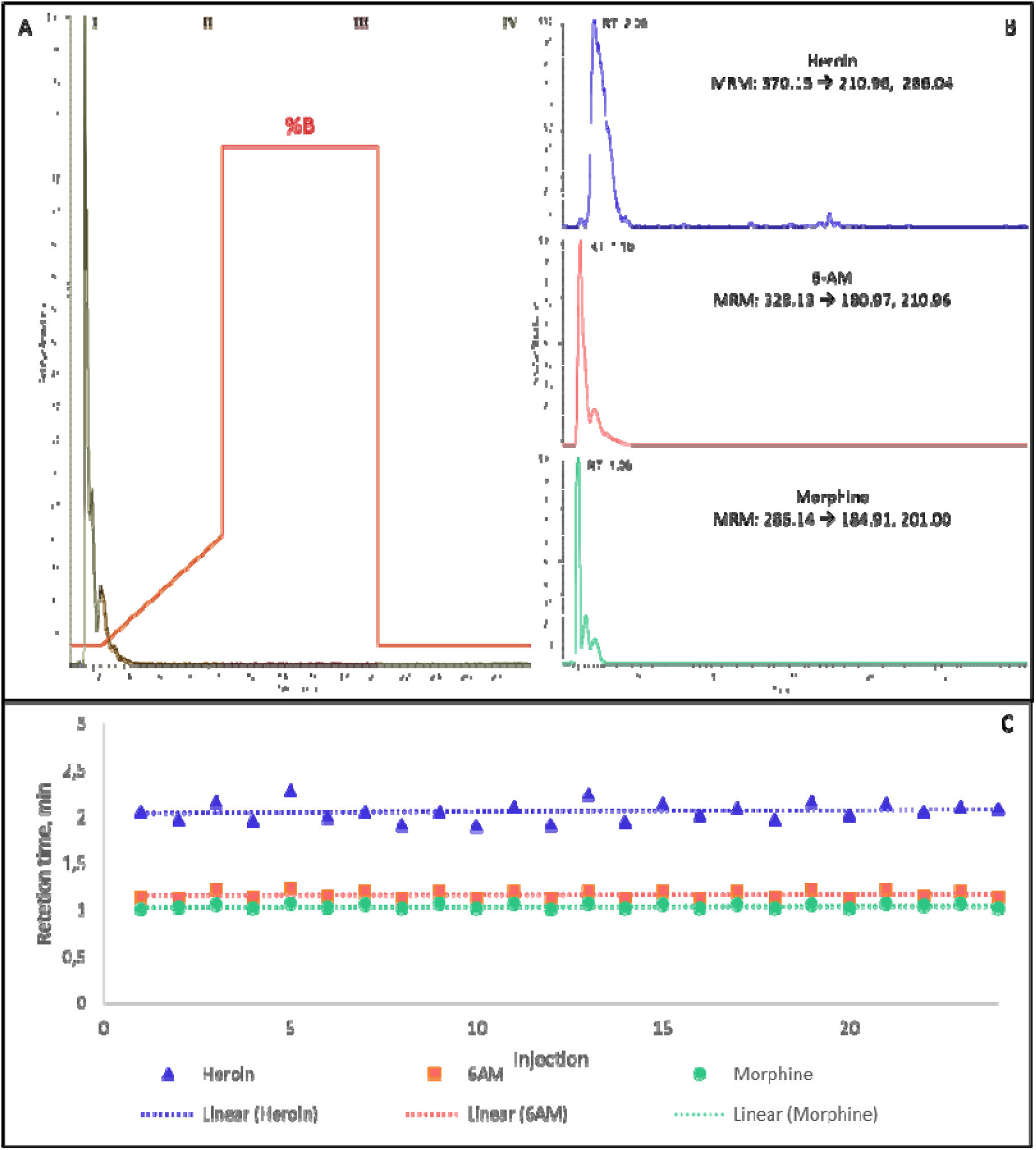
**A:** LC-MS chromatogram of heroin and metabolites derived from an “organ-in-a-column” system exposed to heroin coupled on-line with liquid chromatography-mass spectrometry. The gradient program as a function of organic mobile phase modifier (%B) is shown in red. The four stages of the gradient program are (I) isocratic elution, (II) gradient elution, (III) wash and (IV) re-equilibration. B: Extracted ion chromatograms for heroin and metabolites are shown with MRM-transitions that were used. The chromatograms were extracted from the same run that is shown in figure 5A. C: Retention time variability of heroin and metabolites over an entire exposure experiment (24 injections/12 hours).

### Temperature controlled drug delivery ensures improved robustness

Heroin can spontaneously decompose into its metabolite 6-AM. However, we found that heroin could also be converted to morphine non-enzymatically. At 37 °C, approximately 5% conversion of heroin to morphine was observed after 24 hours and 20% over 120 hours **(see Supporting Information, SI2).** Hence, at physiological temperatures required for metabolic functional cells, heroin can decompose to 6-AM and morphine in the absence of liver organoids. However, when cooled to 4 °C, less than 1% morphine was formed in the absence of liver organoids **(see Supporting Information, SI2).** To avoid formation of morphine prior to organoid exposure, a 3D-printed syringe cooler was designed and implemented in the system (**Figure 1**).

### “Organ-in-a-column”-LC-MS drug metabolism studies on iHLC

Our next step was to evaluate the system’s functionality for tracking drugs and metabolites over time, with cooled organoid medium supplemented with 10 μM heroin and iHLC organoids co-loaded with glass beads. **Figure 6** shows the degradation of heroin and the corresponding generation of metabolites 6-AM and morphine for individual “organ-in-a-column” units, as documented by three experiments performed with iHLC organoids generated from the WTC-11 cell line, on different days and columns.

Controls (i.e. columns loaded with beads but not loaded with organoids) generated non-detectable amounts of morphine during the 12-hour experiment, establishing that the pre-column cooling system efficiently prevented non-enzymatic degradation. As expected, a gradual conversion of heroin to 6-AM was detected in columns without organoids. In contrast, in organoid-loaded columns heroin levels decreased more rapidly, while morphine levels increased over time. The data presented in Figure 6 were generated over 12 hours of continuous and fully automated/unsupervised analysis, suggesting that on-line coupling of organoids and MS via commercial LC hardware is feasible.

**Figure 6:**
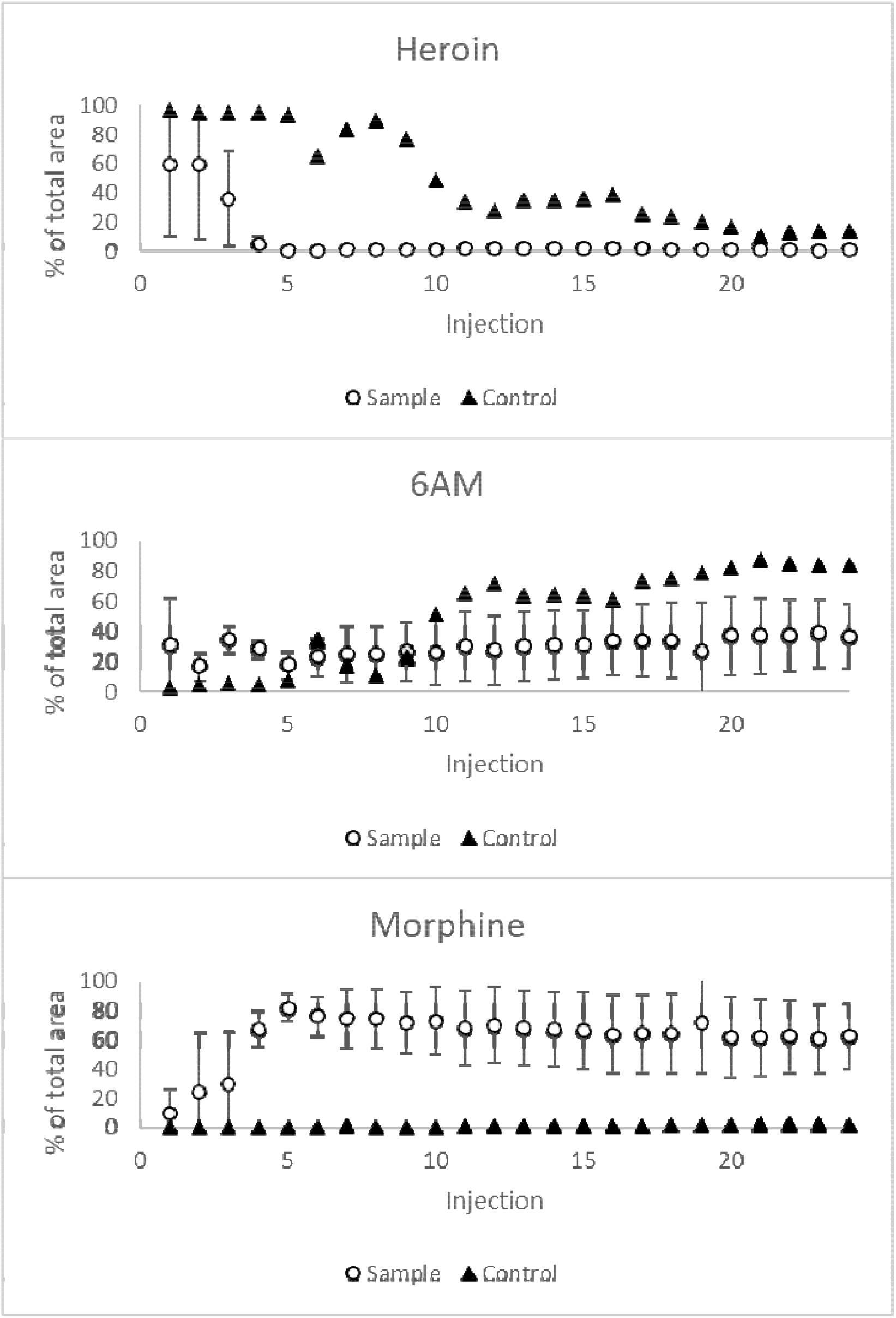
On-line enzymatic and non-enzymatic conversion of heroin (10 μM) to 6-AM and morphine in columns containing approximately 50 iHLC organoids (sample) and columns containing no organoids (control). Graphs show average areas of heroin and metabolites normalized to the average total area of heroin, 6-AM and morphine (area-% of avg. total area of heroin, 6-AM, and morphine). The average total areas of heroin, 6-AM and morphine are based on three runs performed on different days (12 h experiments) and columns with organoids from the WTC-11 cell line (error bars: standard deviation, n=3).

Next, heroin metabolism was compared in columns using iHLC organoids independently differentiated from three iPSCs. As expected, the biological variation in these experiments resulted in larger standard deviation, but still identified significant levels of morphine when compared to the control (n=3) (see **Supporting Information, SI3).**

### Concluding remarks

In this proof-of-concept study, liver organoids have been loaded in liquid chromatography column housings (“organ-in-a-column”) and coupled on-line with mass spectrometry for direct analysis of drug metabolism. Features of the here described setting include a substantial degree of automation compared to our previous manual efforts [10], selective measurements through multiple reaction monitoring, and an increased degree of robustness through the use of standard LC parts and fittings, compared to non-commercial chips previously employed [14]. The setup could be used for directly identifying liver organoid-induced drug metabolism, and subsequent hour-scale monitoring of metabolism. Future qualitative identifications of metabolites will include further standardization of column packing, and inclusion of internal standards to reduce ESI-MS signal variations.

The system will be further explored for additional drugs and configurations (a next step will be to include an autosampler for multi-drug analysis. This encourages next steps to include expanded drug metabolism studies for mapping enzyme activity (e.g. drugs such as phenacetin, tolbutamide and fluoxetine, metabolized by CYP2D6, CYP2C9 and CYP1A2, respectively), which can have clear benefits in e.g. personalized drug development when assessing organoids grown from individual patients/patient groups.

## Supporting Information

Additional experimental details, materials, and methods, regarding live/dead staining (S1), heroin stability tests (S2), proteomics (S3) and mRNA studies (S4).

## Acknowledgments

This work was supported by the Research Council of Norway through its Centre of Excellence scheme, project number 262613, and project number: 295910 (National Network of Advanced Proteomics Infrastructure). Support was also provided by UiO:Life Science and the Olav Thon Foundation.

