## Supporting Information for "“Organ-in-a-column” coupled on-line with liquid chromatography-mass spectrometry"

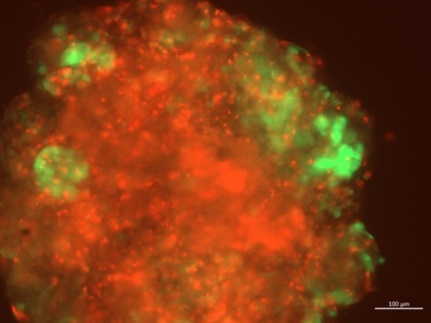

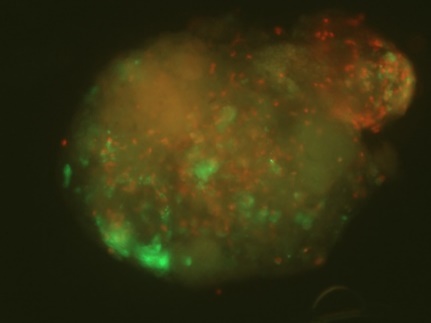

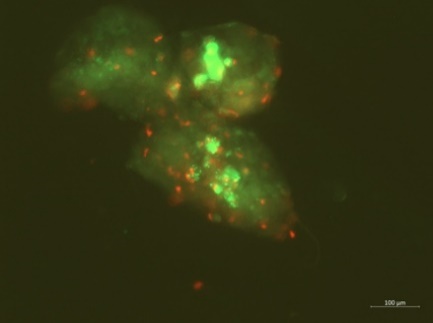

**Figure SI1**. Live/dead staining of liver organoids flushed from the LC column after 7 days on-line experiments. Live cells were stained green. Most (>80%) organoids showed low amounts of dead staining (red color, middle, right) even after exposure to high concentrations of heroin and left under perfusion of medium without oxygenation for more than 7 days. However, some in-column variation was seen with some organoids showing >90% dead stain (left). Scale bar is 100 µm.

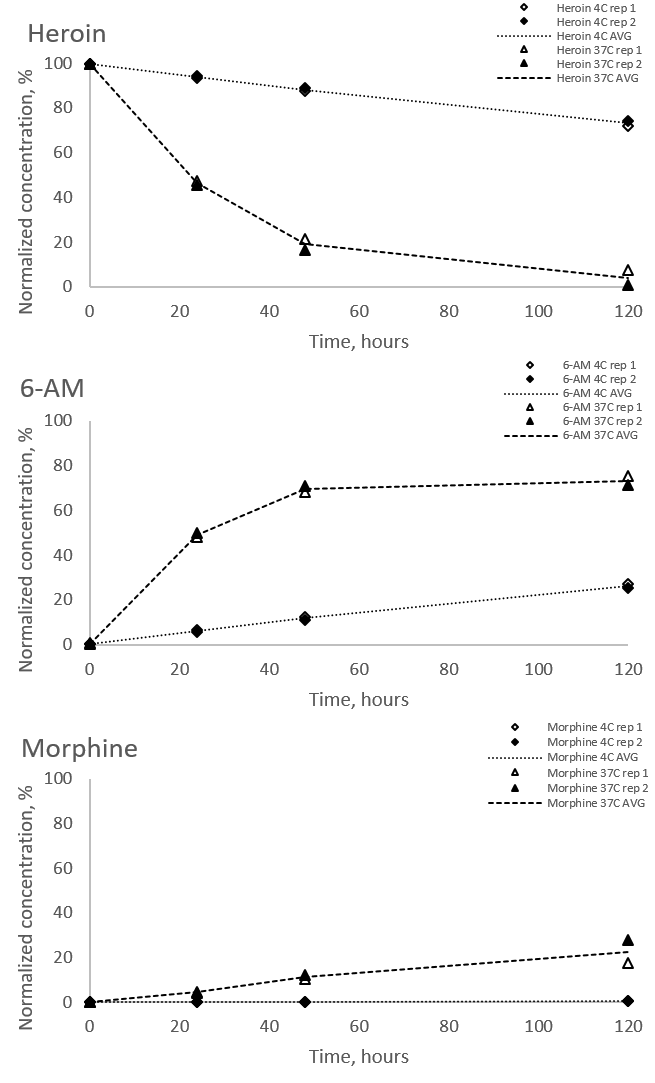

**Figure SI2.** Heroin stability testing: Spontaneous cell-independent degradation of heroin (10 µM) formation of 6-AM and morphine from heroin at different temperatures in serum free organoid medium (4 ˚C vs 37 ˚C).

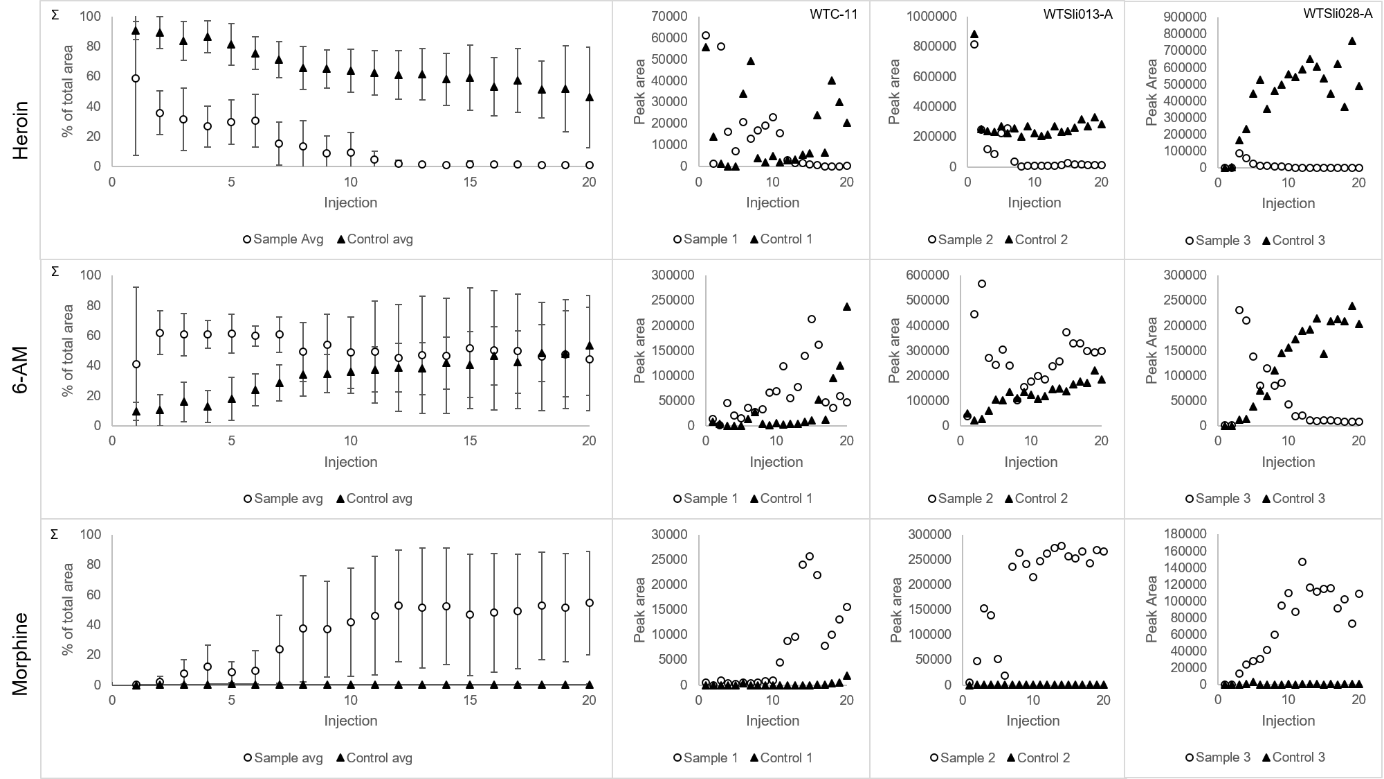

**Figure SI3**. On-line enzymatic and non-enzymatic conversion of heroin (10 µM) to 6-AM and morphine in columns containing approximately 50 organoids (sample) and columns containing no organoids (control). The top row shows average areas of heroin and metabolites normalized to the average total area of heroin, 6-AM and morphine (area-% of avg. total area of heroin, 6-AM, and morphine). The average total areas of heroin, 6-AM and morphine (top row) are based on three experiments performed on different days (10 h experiments) and columns, with three different cell lines displayed with the name of the cell line.

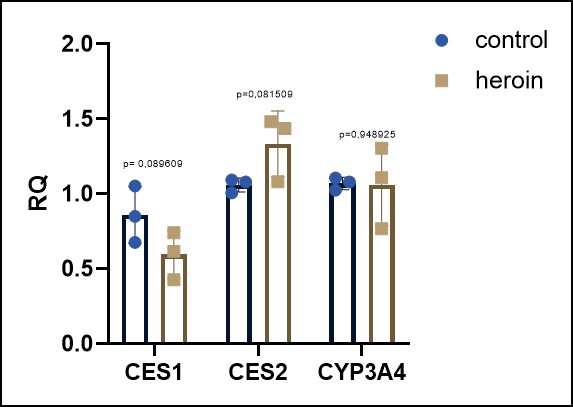

**Figure SI4.** Relative expression of CES1, CES2 and CYP3A4 in heroin-treated and control iHLC organoids.

Significance was calculated using unpaired t-test.

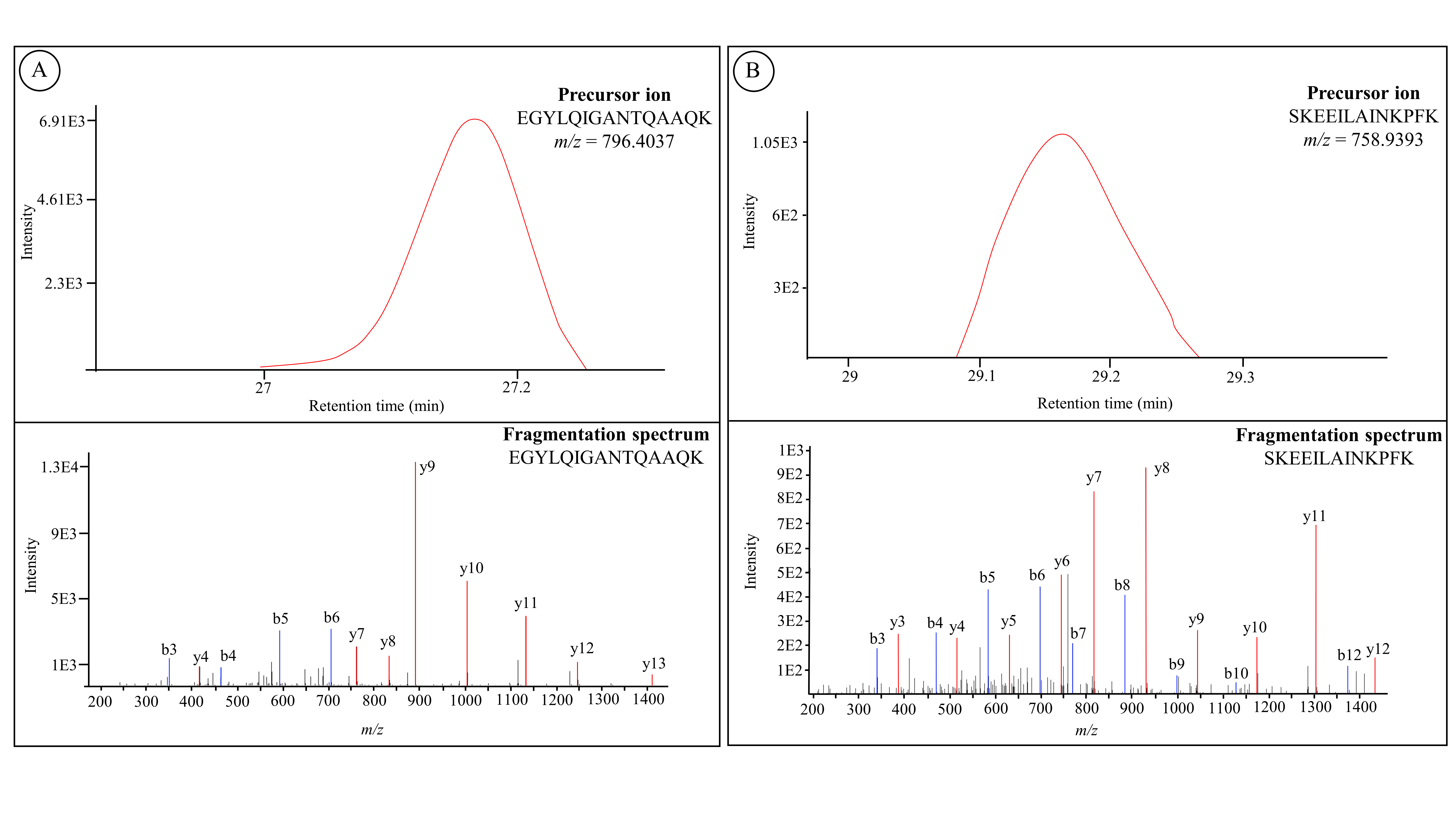

**Figure SI5.** Detection of human liver carboxylesterase 1 (CES1, **A**) and human liver carboxylesterase 2 (CES2, **B**) in liver organoids. The example shown in this figure is iHLC generated from WTC-11 cell line, treated with heroin (10 µM). The figure shows the extracted ion chromatograms (top) of the unique tryptic peptides EGYLQIGANTQAQK (for CES1) and SKEEILAINKPFK (for CES2) and the fragmentation spectrums (bottom) with detected y and b ions marked in red and blue, respectively. The peptides were separated using a 25 cm x 75 µm IonOpticks column (1.6 µm particles). The mobile phases contained 0.1% formic acid in water (A) and 0.1% formic acid in acetonitrile (B). A linear gradient from 0-35% mobile phase B over 54 min at a flow rate of 300 nL/min at a column temperature of 50 °C was employed. MS acquisition was performed using a timsTOF in data-dependent acquisition parallel accumulation-serial fragmentation (DDA-PASEF) mode. Detection was performed with at least 1 unique peptide and a false discovery rate at ≤1%.
